# Interspecific adaptations in root system architecture define host tolerance of Arabidopsis to biotic stresses by root feeding nematodes

**DOI:** 10.64898/2026.04.08.717173

**Authors:** Jaap-Jan Willig, Casper C. van Schaik, Rick Faesen, Sreelekshmi Suresh, Mark G. Sterken, Misghina G. Teklu, Geert Smant

**Affiliations:** Laboratory of Nematology, Wageningen University & Research, 6708PB Wageningen, The Netherlands; Agrosystems Research, Wageningen University & Research, 6708PB Wageningen, The Netherlands

**Keywords:** Root system architecture, plasticity, adventitious lateral roots, gall formation, tissue damage, tolerance, plant-parasitic nematodes, Arabidopsis

## Abstract

Belowground, plants are exposed to a wide range of biotic stresses that vary in severity and nature, including tissue damage, disruption of vascular connectivity, and depletion of assimilates. How plants adapt their root systems to cope with different types of belowground biotic stresses is not well known. In this paper we compare above- and belowground plant adaptations to three nematode species with distinct tissue migration and feeding behaviours to study mechanisms underlying tolerance to different types of biotic stresses. We monitored both green canopy growth and changes in root system architecture of Arabidopsis inoculated with *Pratylenchus penetrans*, *Heterodera schachtii*, and *Meloidogyne incognita*. This revealed three distinct phases in aboveground plant responses: (i) initial growth inhibition associated with host invasion and tissue damage, (ii) persistent growth reduction associated with nematode sedentarism, and (iii) late growth stimulus in more advanced stages of infection. Specific adaptations in the root systems further revealed fundamentally different stress coping strategies. Tissue damage and intermittent feeding by *P. penetrans* in the root cortex did not induce significant changes in root system architecture. Tissue damage to the root cortex and prolonged feeding on host vascular cells by *H. schachtii* induced secondary root formation compensating for primary root growth inhibition. Prolonged feeding on host vascular cell by *M. incognita* alone did not induce secondary root formation, but was accompanied by typical local tissue swelling instead. Our data suggest that local secondary root formation and tissue swelling are two distinct compensatory mechanisms underlying tolerance to sedentarism by root-feeding nematodes.

**Highlight:** How plants utilize root system plasticity to cope with different types of biotic stresses by root feeding nematodes remains largely unknown. Here, we report on specific adaptive growth responses in Arabidopsis roots to three nematode species, *Pratylenchus penetrans*, *Heterodera schachtii*, and *Meloidogyne incognita*, with fundamentally different strategies for host invasion, subsequent migration through host tissue, and feeding on host cells.

## Introduction

Disease tolerance refers to the ability of plants to mitigate the impact of biotic stress by pathogens and herbivorous attackers (Peterson et al., 2017; Willig et al., 2023). Conceptually, tolerance contrasts with resistance, which focusses on reducing the level of biotic stress by targeting the causal agent. Many studies on the genetic and molecular mechanisms governing disease resistance have been published over the past forty years (Jones et al., 2024). However, far less is understood of the physiological processes underlying disease tolerance in plants. Nonetheless, disease tolerance is thought to be a complex plant trait, because biotic stresses are also often complex in nature. For instance, biotic stress in plants by sedentary root-feeding nematodes is a convolution of tissue damage in the root cortex and root apical meristem, reduced uptake and transport of water and minerals from the soil, and depletion of plant assimilates for root growth, all of which do not necessarily occur at the same time and place. This means that the plant responses mitigating the impact of root feeding by nematodes are most likely multilayered and diverse in time and space as well.

The microscopically small root-feeding nematodes form a useful model for studying tolerance to belowground biotic stress in plants, because of their diversity in host invasion and feeding strategies (Goverse & Smant, 2014; Hofmann & Grundler, 2007). Individual nematodes can take weeks to develop into the reproductive stage, whilst all the time feeding on living host cells, sometimes even without moving about (Wyss, 1992; Wyss & Grundler, 1992). The most basal nematode lineages feed on different host cells from outside plants and remain mobile throughout all development stages (Perry & Moens, 2011; Quist et al., 2015). Endoparasitic lineages feed on host cells from within the host, some of which also remain mobile throughout the entire lifecycle (Zunke, 1990b), while others become sedentary shortly after establishing a permanent feeding site (Wyss, 1992; Wyss & Grundler, 1992). Migratory endoparasites extract nutrients from multiple living host cells and migrate intermittently inside host tissue between feeding periods (Zunke, 1990a, 1990b). The most advanced sedentary endoparasitic nematodes also migrate during the early stages of infection, but only to find a suitable place in the root to establish a permanent feeding site, through which they gain access to the flow of assimilates in the plant’s vascular system. Nematode migration can be intracellular, resulting in a trail of dead cells along the migratory tract (Rehman et al., 2009). However, some nematode species move about inside plant tissue by enzymatically weaking the middle lamella and pushing host cells aside, causing little overt damage (Grundler et al., 1994; Zhou et al., 2019). Thus, a differential analysis of major lineages of root-feeding nematodes may offer opportunities to resolve the molecular and genetic mechanisms underlying tolerance to a whole range of specific biotic stresses (e.g., loss of cellular integrity, disruption of vascular connectivity, and depletion of resources).

Recently, we used aboveground plant growth responses of *Arabidopsis thaliana* to study differences in tolerance to root-knot nematodes (i.e., *Meloidogyne incognita*) and cyst nematodes (i.e., *Heterodera schachtii*) (Willig et al., 2023). Members of both families are sedentary endoparasites which draw their nutrients from host phloem and xylem via a permanent feeding site, but they do so differently. Infective juveniles of root-knot nematodes migrate intercellularly through host tissue in the cortex to the root apical meristem where they transform undifferentiated vascular parenchyma cells just above the cell division zone into permanent feeding cells, so-called giant cells (Kyndt et al., 2013; Olmo et al., 2020). Individual root-knot nematodes typically induce a few giant cells, each of which provides the feeding nematode access to neighbouring phloem and xylem elements. By contrast, infective juveniles of cyst nematodes migrate intracellularly through the root cortex for a few hours after which they transform a host cell located in the cortex, endodermis, or pericycle into an initial syncytial cell (Wyss & Zunke, 1986). Feeding on this host cell triggers progressive, local plant cell wall degradation and subsequent cell fusion with neighbouring vascular cells (Bohlmann & Sobczak, 2014), which ultimately results into the incorporation of hundreds of vascular parenchyma cells into a syncytium closely connected to phloem and xylem elements (Wyss, 1992). Neither root-knot nematodes nor cyst nematodes move about in between feeding periods, but in other aspects their differences may be causal to differences in tolerance of Arabidopsis to biotic stress by these worms.

To quantify differences in tolerance to infections by root-knot nematodes and cyst nematodes, we built a high throughput phenotyping platform using the green canopy area of Arabidopsis plants as a proxy for belowground adaptations to nematode infections (Willig et al., 2023). Quantifying such differences in tolerance by means of a single metric is challenging. From an agronomic perspective the minimal yield by population density of an attacker is most relevant (Seinhorst, 1986; Teklu et al., 2022). However, this is an endpoint measurement which does not help resolving the underlying tolerance mechanisms during earlier stages of the infection cycle. To address this shortcoming, Seinhorst (1986) coined the term tolerance limit (*Tl*) for the lowest population density of nematodes at which a significant decline in a plant growth parameter can be observed at a given point in time (Seinhorst, 1986). However, Seinhorst’s approach relies on how well the observations fit an inverted sigmoid model, which disregards informative variation in output at lower population densities. To take this variation into account, we developed a novel analytical approach wherein observations on plant growth parameters (e.g., green canopy area) are used to estimate the maximum growth output (*K*) and the intrinsic growth rate (*r*) (i.e., maximum achieved growth rate irrespective of time (Willig et al., 2023)). Altogether, the tolerance limits, the maximum growth outputs, and intrinsic growth rates revealed remarkable differences in how plants mitigate the impact of biotic stress by root-knot nematodes and cyst nematodes.

How plants cope with stress induced by endoparasitic nematode lineages that do not form a permanent feeding site inside the host and thus remain migratory throughout their lifecycle is not understood. To address this knowledge gap, we focused on the root-lesion nematode *Pratylenchus penetrans*, which is a migratory endoparasite of many plant species (Castillo & Vovlas, 2007). *Pratylenchus penetrans* invades a host plant at the root elongation zone after which it migrates intracellularly through the cortex (Zunke, 1990a, 1990b). Based on observations with different plant species, it is thought that *P. penetrans* exclusively remains in the root cortex (Duncan & Moens, 2013), where it alternates migration with brief and extended feeding periods. Brief feeding lasts only a few minutes and does not involve major structural changes in host cells (Zunke, 1990b). By contrast, extended feeding on single host cells can last for several hours and is always associated with clearly observable subcellular changes. Invariably, extended feeding periods end with host cell death, resulting in further damage to the root cortex, ultimately leading to large lesions consisting of dead epidermal and cortical tissue.

Although *P. penetrans* has a wide host range, little is known of its behaviour and reproduction in Arabidopsis and it was therefore not clear if our tolerance-phenotyping platform could be used for this nematode species (Dinh et al., 2014; Wyss & Grundler, 1992). We therefore first set out to investigate if *P. penetrans* can feed and reproduce on Arabidopsis in culture media and soil. Next, we assessed if stress by *P. penetrans* on the root system of Arabidopsis affects above-ground plant development and growth. Hereto, we inoculated Arabidopsis seedlings with increasing densities of *P. penetrans* and measured the green canopy area, plant height, the number of branches, and the number of flowers for three weeks after inoculation (Willig et al., 2023). Based on this data we subsequently determined the tolerance limit, the maximum growth output, and intrinsic growth rate of Arabidopsis seedlings infected with *P. penetrans*, allowing for a three-way interspecific comparison with *M. incognita* and *H. schachtii*. As aboveground plant responses to root-feeding nematodes often reflect belowground adaptations in root system architecture, we also monitored primary root growth, secondary root formation, secondary root growth, and total root growth by inoculation density and over time. We used this data to link specific compensatory root plasticity mechanisms to tolerance to different types of biotic stress by root-feeding nematodes.

## Materials and Methods

### Plant culturing

For plant growth experiments on culture media, *A. thaliana* Col-0, or *arf7-1/arf19-1* seeds were vapor sterilized for 3-4 hours using a mixture of hydrochloric acid (25%) and sodium hypochlorite (50 g/L). Finally, sterile seeds were stratified for 4 days at 4°C, after which they were sown on six-well or square (120 mm x 120 mm) petri dishes containing either sand, Gamborg B5 media (0.328% Gamborg B5, 2% sucrose, 1.5% Bacto-agar, pH 6.2), modified KNOP media (Sijmons et al., 1991), or Murashige and Skoog (MS) medium with vitamins (4.7 g/l, Duchefa Biochemie), 2% sucrose, and Gelrite (5 g/l, Duchefa Biochemie). Plants were incubated in a growth cabinet with a 16 h-light/8 h-dark photoperiod at 21°C.

For plant growth experiments on sand, non-sterilized Arabidopsis seeds (Col-0) were first stratified for four days at 4°C and then sowed directly on top of on a layer of silver sand in pots in our high-throughput phenotyping platform as described previously (Willig et al., 2023). Seedlings were incubated at 21°C and 16-h-light/8-h-dark photoperiod in a climate room fitted with LED light (150 lumen).

For propagation of *Pratylenchus penetrans*, maize (*Zea mays*) seeds were stratified for four days at 4°C and sowed in 6-liter pots containing in layers from bottom to top: Ederol filter paper (No. 261, J.H. Ritmeester, Utrecht), hydro grains (Lecanto 2-5 mm; bag á 50 l = 25 kg; firm Gebr. Eveleens), silver sand, and perforated black plastic cover (Brinkman’s-Gravenzande Holding). Maize plants were maintained in a greenhouse at 18-22°C, 60-70% relative humidity, and 16h-light/8 h-dark conditions.

### Preparing inoculum of *Pratylenchus penetrans*

Two-week-old maize plants were inoculated with mixed stages of *P. penetrans* (Drenthe population, The Netherlands) using a pipette Stepper in 5 cm deep holes made by a spoon around the periphery of the seedlings (*P_i_* = 4 individuals per gram sand. For a period of 90 days, the sand in the pots was kept moist by adding water including basic plant nutrients. Thereafter, the shoots were cut, while the roots were collected by sieving and washing the sand through 3 mm mesh sieves. Next, the roots were cut into pieces of 1 cm and placed on extraction sieves (150 μm mesh; *D* = 20 cm). Extraction sieves were subsequently placed on top of an extraction dish (*D* = 25 cm) and incubated in a mist chamber (Seinhorst, 1988). Nematode suspensions harbouring mixed stages were collected every 3 days from the mist chamber and stored at 4°C until the time of inoculation.

Nematode suspensions were further purified by centrifugation on a 35% sucrose gradient, transferred to a 2 mL Eppendorf tube, and surface sterilized for 15 minutes in a solution containing 0.16 mM HgCl_2_, 0.49 mM NaN_3_, and 0.002% (v/v) Triton X-100. After washing the nematodes three times with sterile tap water, they were re-suspended in a sterile 0.7% Gelrite (Duchefa Biochemie, Haarlem, the Netherland) solution to be used as inoculum in further experiments. A similar concentration of Gelrite solution lacking nematodes was used as mock inoculum.

### Monitoring *Pratylenchus penetrans* infection in Arabidopsis roots *in vitro*

Sterile Arabidopsis seeds were incubated for four days on solid Gamborg B5 media in square Petri dishes (12 cm), after which they were inoculated with 70 surface-sterilized *P. penetrans* (mixed stages: J2s to adult males and females). The infection process was monitored for a period of 24 days. Images or video recordings of nematode infected roots were made with an SZX10 Stereomicroscope (Olympus) or Axio Imager.M2 LED transmitted light microscope (Zeiss). Images of roots were made using a Perfection V800 Photo flatbed scanner (Epson, Nagano, Japan).

### Quantifying reproduction of *P. penetrans* in Arabidopsis in sand

Briefly, prior to sowing, pots were filled with dry silver sand, covered with black coversheets, and incubated in Hyponex (1.7 mM/L NH_4_^+^, 4.1 mM/L K^+^, 2 mM/L Ca_2_^+^, 1.2 mM/L Mg_2_^+^, 4.3 mM/L NO_3_^-^, 3.3 mM/L SO_4_^2-^, 1.3 mM/L H2PO_4_^-^, 3.4 µm/L Mn, 4.7 µm/L Zn, B 14 µm/L, 6.9 µm/L Cu, 0.5 µm/L Mo, 21 µm/L Fe, pH 5.8) for five minutes. Seven days after sowing, seedlings were incubated again for five minutes in Hyponex. Nine-day-old seedlings were inoculated with increasing densities of *P. penetrans* (0 to 100 nematodes per g dry sand; mixed stage inoculum) and grown for a period of 35 days. Next, nematode densities from the root fraction and sand fraction of each pot were determined. The roots of each pot were collected by first gently sieving through a 3-mm mesh sieve. Subsequently, roots were transferred to an extraction dish of 6 cm diameter and placed in a mist-chamber (Seinhorst, 1988). Suspensions of nematodes were tapped off every week for 28 days. Nematodes that remained in the sand fraction were extracted using Seinhorst elutriators. Next, using the Seinhorst population dynamics model (equation 1) the final population density was described:

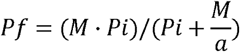

Where *P_f_* is the final population density, *P_i_* is the initial population density, *a* is the maximum multiplication rate, and *M* is the maximum population density. The relationship between *P_i_* and *P_f_* values was visualized, and the population dynamics model was fitted to the mean *P_f_* values using parameters obtained through non-linear regression analysis. The Hessian matrix was used to estimate standard errors and 95% confidence intervals for the parameters, and model fit was assessed using the adjusted *R*², accounting for degrees of freedom.

### Analysing green canopy area of nematode-infected Arabidopsis plants

The green canopy of individual nematode-infected Arabidopsis plants was captured and analyzed as previously described (Willig et al., 2023). Briefly, every hour, pictures were taken of the plants (15 pictures per day) for a period of 21 days. At the end of the experiment, colour corrections were done using Adobe Photoshop (Version: 22.5.6 20220204.r.749 810e0a0 x64). The surface area of the rosette was determined using a custom-written ImageJ macro (ImageJ 1.51f; Java 1.8.0_321 [32-bit]) and Java was used to make GIFs.

### Modelling plant growth and tolerance using a high-throughput phenotyping platform

Data of aboveground plant growth of *P. penetrans* infected Arabidopsis seedlings was analysed as described previously (Willig et al., 2023); available via Gitlab: https://git.wur.nl/published_papers/willig_2023_camera-setup). For comparison, we retrieved data of aboveground plant responses of Arabidopsis seedlings inoculated with *H. schachtii* and *M. incognita* using the exact same set up from a previous study (Willig et al., 2023).

In short, the measurement used was the median green canopy area (cm^2^), calculated from the 15 daily measurements. We used log_2_-transformed data, where the rate of growth was determined per day per plant by (equation 2)

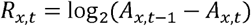

where *R_x,t_* is the transformed growth rate of plant *x* at day *t* from day *t-1* to day *t* based on the median green canopy area *Ax,t*.

The tolerance limit was estimated by fitting the data on a logistic growth model (Willig et al., 2023) using the *growthrates* package on the median daily green canopy area A_t_ (cm^2^) (equation 3),

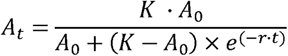

wherein *K* is the maximum green canopy area (cm^2^), *A_0_* is the initial canopy area (cm^2^), and *r* is the intrinsic growth rate (d^-1^), which were determined as a function of time *t* (d) (p < 0.1).

Based on the relation between *K* and inoculation density we could identify the tolerance limit (equation 4)

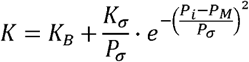

where *P_i_* is the initial nematode density in nematodes per gram dry sand, *K_B_* is the green canopy area size, *K_σ_* is the normalized maximum green canopy area that can be achieved over the *P_i_* range, *P_σ_* is the deviation around the nematode density allowing maximum growth, *P_M_* is the nematode density at which maximum growth is achieved. We modelled the parameter values using *nls* and extracted confidence intervals using the *nlstools* package (Baty et al., 2015). The tolerance limit, 2\**P_M_,* was determined as in (Willig et al., 2023).

### Hatching of *H. schachtii* and *M. incognita* and sterilization of nematodes

Cysts of *H. schachtii* (Woensdrecht population from IRS, The Netherlands) were collected from the sand and roots of infected *Brassica oleracea* plants following the method described by Baum *et al*., 2000. Eggs of *M. incognita* (strain Morelos from INRA, Sophia Antipolis, France) were extracted by soaking nematode-infected roots in 0.05% (v/v) NaOCl for 3 mins. The roots were then rinsed with tap water, and the eggs were collected using a 25µm sieve. Subsequently, *M. incognita* eggs and *H. schachtii* cysts were using 0.02% of NaN_3_ and kept for 20 mins undisturbed. Then, the cysts and eggs were thoroughly washed in tap-water to remove all traces of NaN_3_ and transferred to a hatching sieve. Finally, the sieve was placed in a glass petri dish containing an antibiotic solution of 1.5 mg/ml gentamycin, 0.05 mg/ml nystatin in water, and 3mM ZnCl_2_ for *M. incognita* eggs, and *H. schachtii* cysts, respectively. Both the cysts and eggs were incubated in the dark room at room temperature for 4-7 days for *H. schachtii* and *M. incognita*. For sterile experiments with *P. penetrans*, juveniles were collected as described above but incubated for 4-7 days in the same antibiotic solution as *M. incognita*.

After hatching of *H. schachtii* and *M. incognita*, and collection of *P. penetrans*, juveniles were transferred to a conical glass centrifuge tube and the purification was done with equal volume of 70% sucrose. After washing three times by centrifugation, the nematodes were suspended in 2 ml Eppendorf tube containing Tween-20 solution and water. Later, the surface sterilization of juveniles were done with 0.4% Triton x-100, 1.0% NaN_3_ and 0.8% HgCl_2_ for 15 mins for *M. incognita* and 20 mins for *H. schachtii* and *P. penetrans*. After incubation, the juveniles were washed thoroughly three times with sterile water, removing the supernatant each time. Finally, the juveniles were resuspended in 0.7% gelrite.

### Quantifying root system architecture of nematode-infected Arabidopsis

Nine-day-old Arabidopsis seedlings, grown on 120×120 mm square Petri dishes were inoculated with 0 (mock), 0.5, 1.0, 2.5, 5.0, 7.5, 10, 15, 20, and 25 juveniles per milliliter of modified Knop medium as previously described (Willig et al., 2024). Inoculations were done with two 5-µL drop that were pipetted at opposite sides of each seedling while keeping the Petri dishes vertical. Seedlings were grown vertically and covered by a light-blocking paper for seven days. At 7 dpi, scans were made of whole seedlings using an Epson Perfection V800 photo scanner. The architecture (i.e. total root length, primary root length, total secondary root length, the number of root tips, and average secondary root length) were measured using SmartRoot (Lobet et al., 2011) and ImageJ version 1.53.

### Statistical analysis

Statistical analyses were performed using the R software version 3.6.3 (Windows, x64). The R packages used are *tidyverse* (https://CRAN.R-project.org/package=tidyverse), *ARTool* (https://CRAN.R-project.org/package=ARTool) and *multcompView* (https://CRAN.R-project.org/package=multcompView). Correlation between variables was calculated using Spearman Rank-Order Correlation coefficient. For normally distributed data, significance of the differences among means was calculated by ANOVA followed by Tukey’s HSD test for multiple comparisons. A non-parametric pairwise Wilcoxon test followed by false discovery rate correction for multiple comparisons was used for data with other distributions and one grouping factor. For the high-throughput platform data we used the Wilcoxon test as implemented in the *ggpubr* package (https://cran.r-project.org/web/packages/ggpubr/index.html). The confidence interval of the inoculum density-response curves was calculated by LOESS regression (as per default in *geom_smooth*) in R.

## Results

### Establishing Arabidopsis as a model host for migratory endoparasites

To determine whether the migratory endoparasite *P. penetrans* can invade, feed, and produce offspring in Arabidopsis, we inoculated a mixture of different nematode stages to four-day old *in vitro* cultured Arabidopsis seedlings (Fig. 1a). One day post inoculation (dpi), we observed different life stages of *P. penetrans* invading the roots of Arabidopsis using their oral stylet to puncture plant cell walls (Fig. 1b, Video S1). At 2 and 3 dpi, we found *P. penetrans* moving about inside the root cortex causing extensive damage to root tissue (indicated by browning) (Video S2). We also observed individuals protruding their stylet inside host cells for longer periods of time, which we interpreted as feeding behavior (Video S3). Around 10 dpi, we found *P. penetrans* laying eggs inside the Arabidopsis roots by using fierce tail movements and body contractions (Video S4). At later timepoints, damage to the roots resulted in expansive brown lesions. Most likely this was due to continuing migration and feeding. Based on our observations, we conclude that *P. penetrans* can establish an infection, feed on host cells, and lay eggs in Arabidopsis *in vitro*.

**Figure 1:**
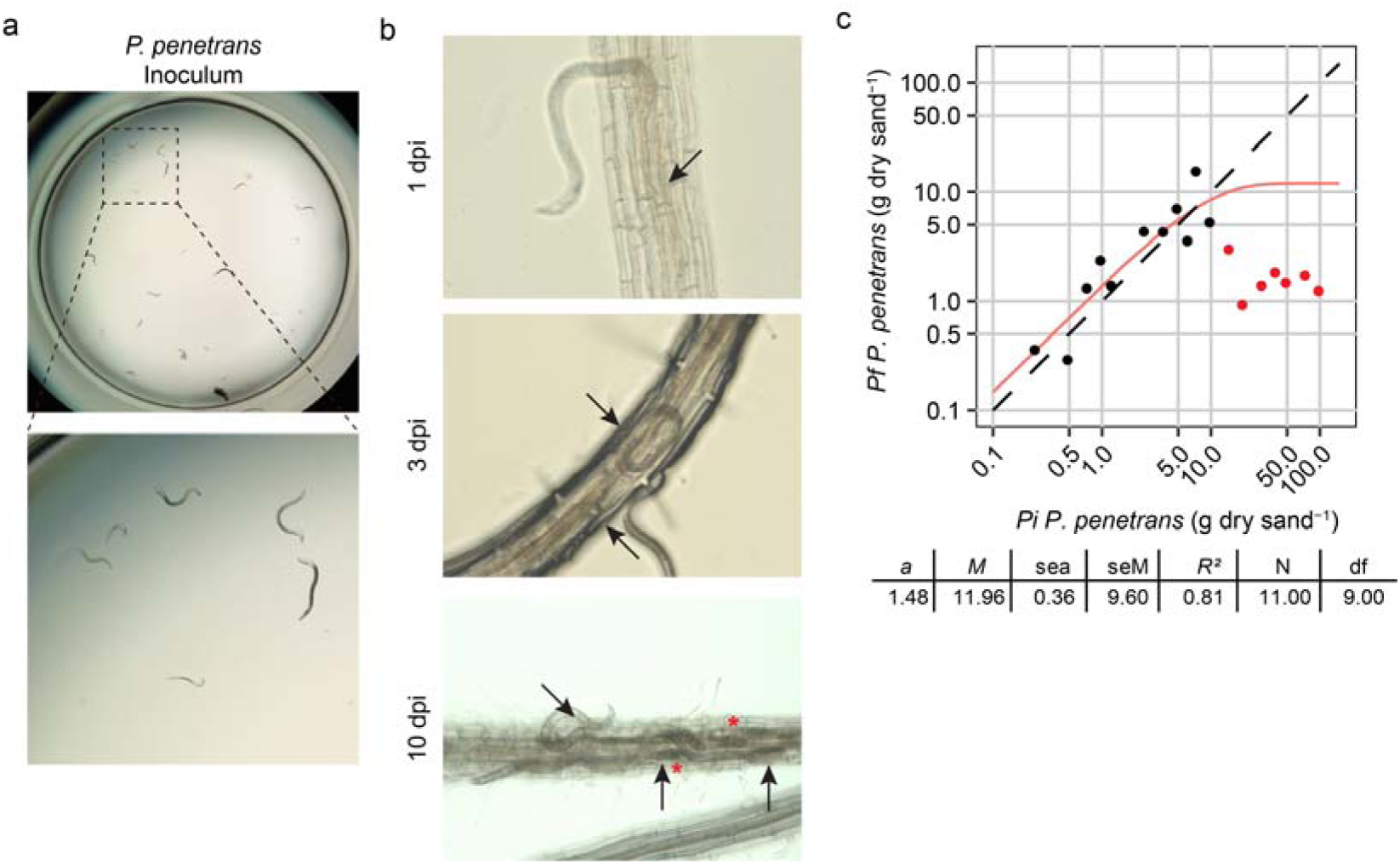
*Pratylenchus penetrans* infection in Arabidopsis roots. Four-day old Arabidopsis seedlings were inoculated with 70 mixed stage juveniles *P. penetrans*. **a)** Inoculum of *P. penetrans*. Sizes indicate different developmental stages of *P. penetrans*. **b)** Infection of *P. penetrans* at 1, 3, and 10 dpi. Arrowhead indicates the head of the nematode. Red asterisks indicate location of eggs. **c)** The quantitative relationship between *P_i_* and *P_f_* of *P. penetrans* in Arabidopsis. Dots indicated the mean *P_f_* value. The solid red line is the fitted line to the model (equation 1). The dotted line is the equilibrium state for the population (*P_i_* = *P_f_*). *a* is the maximum multiplication rate, *M* is the maximum population density, sea is the standard error of a, se*M* is the standard of *M*, *R*^2^ is the goodness of the fit, N is the number of densities included in the fitting of the model, df refers to the degrees of freedom in the analysis. Red dots are excluded from the model.

Our tolerance phenotyping platform measures aboveground plant performance of Arabidopsis seedlings cultivated in sandy soil. To assess if Arabidopsis can stably sustain nematode population levels of *P. penetrans* under these conditions, we determined nematode reproduction at 35 days after inoculating emerging seedlings in sand soil with increasing densities of mixed life stages (Fig. 1c). We excluded the final population density values after inoculating 15 or more *P. penetrans* per g dry sand from our calculations, because above this inoculation density the plants completely collapsed/died (Fig. S2). First, we observed that the final population density (*P_f_*) of *P. penetrans* on Arabidopsis follows the initial inoculation density (*P_i_*) in a typical sigmoid curve with a saturation point close to an initial inoculation density of 10 *P. penetrans* per gram dry sand. Using a logistic model, which explained 81% of the variation in our data, we estimated the maximum multiplicate rate (a) of *P. penetrans* on Arabidopsis at 1.48 and a maximum population density (*M*) of 11.96 *P. penetrans* (per g dry sand).. Based on these results, we conclude that *P. penetrans* can establish an infection and reproduce on Arabidopsis in sandy soil, but it is only able to sustain relatively low population levels on this plant species.

### *P. penetrans* affects Arabidopsis green canopy area in a density-dependent manner

To quantify the impact *P. penetrans* on overall plant performance of Arabidopsis, we monitored aboveground plant growth and other developmental phenotypes of Arabidopsis seedlings inoculated with increasing densities of mixed-stage *P. penetrans* suspensions in sand over a period of 21 days (Fig. 2). In a striking contrast with previous observations from experiments with *H. schachtii* and *M. incognita* (Willig et al, 2023), *P. penetrans* infections resulted in a complete collapse of Arabidopsis at higher inoculation densities (Fig. S2). No aboveground plant growth and development were observed from 5 dpi onwards for inoculation densities above 15 *P. penetrans* per gram of dry sand (Fig. 2a-d). Below this tipping point, the green canopy area of Arabidopsis offered the highest resolution as a response parameter to increasing nematode inoculation densities (Fig. 2a) compared to the other aboveground plant phenotypes (Fig. 2b-d). Already at 1 dpi, the growth rate of the green canopy area of plants inoculated with 0.25 nematodes per gram of dry sand diverged significantly from mock-inoculated plants (Fig. S1). For the following 9 days, the growth rate of the green canopy area of nematode- and mock-inoculated plants diverged further by inoculation density up to 10 nematodes per gram of dry sand. Thereafter, the green canopy growth of these plants (P_i_ < 10) returned to almost the same level as mock-inoculated plants (Fig. S1; days 10 to 21). We therefore concluded that at permissive inoculation densities *P. penetrans* impacts growth and development of Arabidopsis the most during the early stages following exposure, whereafter the plants seem to recover.

**Figure 2:**
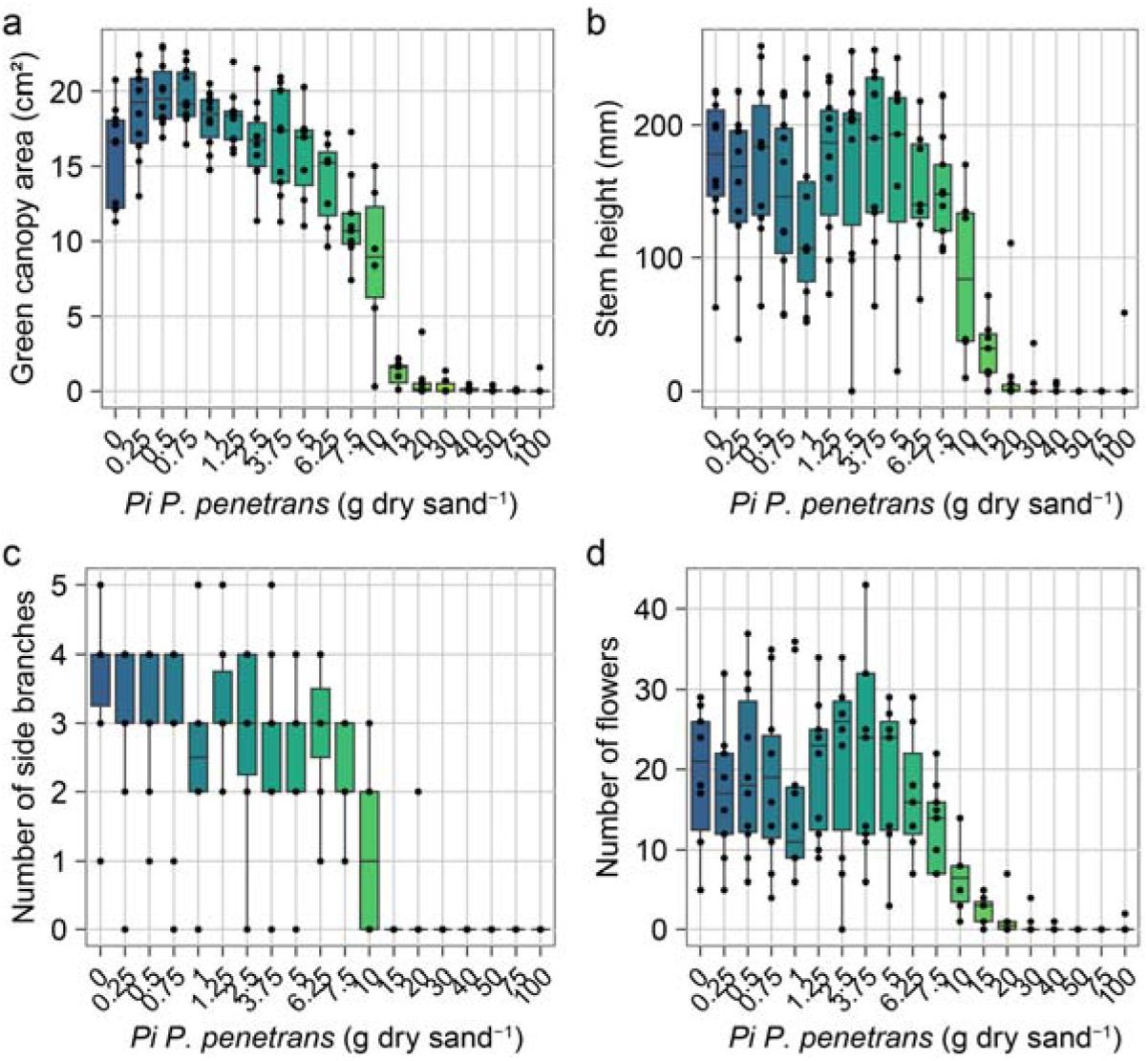
Aboveground growth and development of Arabidopsis respond in a density-dependent manner to *Pratylenchus penetrans* infection. Nine-day old Arabidopsis seedlings were inoculated with 20 densities (*P_i_*) of *P. penetrans* (0 to 100, *P. penetrans* (g dry sand)^-1^ and were monitored for a period of 21 days. Data on **a)** green canopy area, **b)** stem height in cm, **c)** number of side branches, and **d)** number of flowers were collected at 28 dpi. Dots represent individual plants.

### Interspecific differences in tolerance of Arabidopsis to biotic stress by nematodes

Previously, we identified remarkable differences in the growth of the green canopy area and tolerance limit of Arabidopsis infected by *H. schachtii* and *M. incognita* (Willig et al., 2023). To enable a further interspecific comparison between *P. penetrans* and these two sedentary endoparasitic species, we determined the maximum green canopy area (*K*) by inoculation density and derived from that parameter the tolerance limit (*Tl*) of Arabidopsis for *P. penetrans* infections using a logistic growth model (Fig. 3a). The maximum green canopy area of Arabidopsis by increasing inoculation density of *P. penetrans* followed a typical inverted sigmoid curve. Based on the best fitting model (*R^2^*=0.90), the tolerance limit of the Arabidopsis for *P. penetrans* was estimated at an inoculation density of 3.5 *P. penetrans* per gram of dry sand (95% CI: 2.3-4.9). When comparing this outcome with previously published data of *H. schachtii* and *M. incognita* (Fig. 3b and c; Willig et al., 2023), two remarkable differences stand out. First, at low inoculation densities, Arabidopsis infected with *P. penetrans* did not show an enhanced growth response compared to mock-inoculated plants, while this growth stimulus has consistently been observed for both *H. schachtii* and *M. incognita*. Second, the tolerance limits of Arabidopsis for *P. penetrans* and *H. schachtii* (*P_i_* = 3.5; 95% CI: 3.3-3.7) are almost identical, while this is significantly higher for *M. incognita* (*P_i_* = 5.0; 95% CI: 4.3-5.7). Based on these findings, we concluded that the classical bifurcation in feeding behaviours into sedentary vs non-sedentary does not align well with differences in plant tolerance to nematode infection in Arabidopsis. It is, therefore, likely that other factors, such as the level of tissue damage following nematode migration in host tissue during the onset of parasitism, are at play as well.

**Figure 3:**
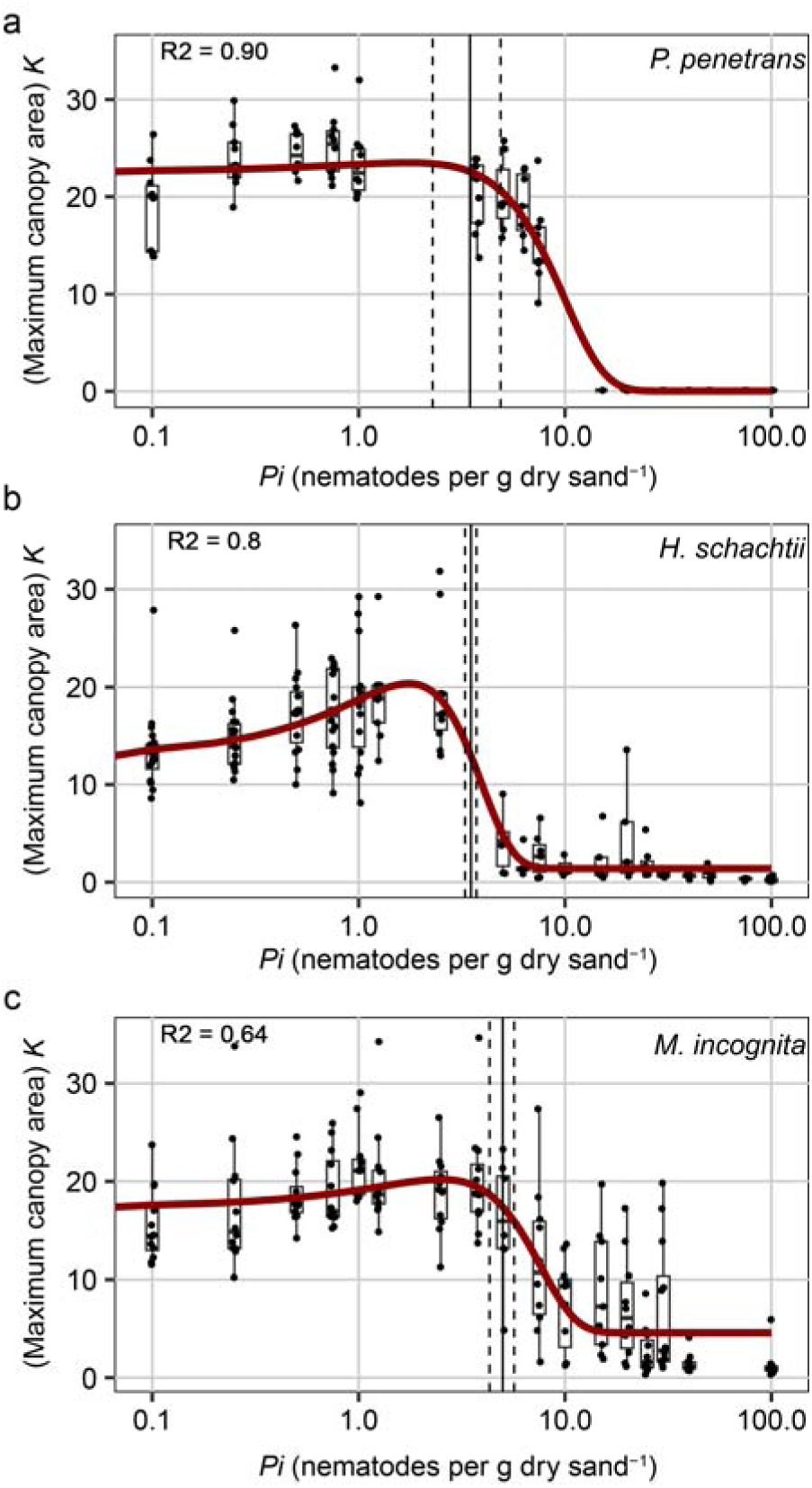
Interspecific differences in maximum green canopy areas of Arabidopsis induced by *P. penetrans*, *H. schachtii*, and *M. incognita*. Nine-day-old Arabidopsis seedlings were inoculated with 18-20 nematode densities of *P. penetrans, H. schachtii*, or *M. incognita* (*Mi*) juveniles (0 to 100 juveniles g dry sand)^-1^. The green canopy area was imaged for a period of 21 days and fitted to a logistic growth rate model (equation 3 and 4). **a)** The maximum canopy area K per inoculation density of *P. penetrans*, **b)** *H. schachtii*, and **c)** *M. incognita*). The fitted line is from a Gaussian curve). Dashed line indicates the confidence interval. *H. schachtii* and *M. incognita* data originate from (Willig et al., 2023).

### Three distinct phases in aboveground plant growth response to nematode infections

*Pratylenchus penetrans* migrates intracellularly throughout its entire lifecycle, resulting in expanding lesions in the root cortex. *Heterodera schachtii* also migrates intracellularly through the root cortex causing significant but limited tissue damage only in the first few days after inoculation. *Meloidogyne incognita* also migrates through the root cortex and root apical meristem in the first few days after inoculation but does not cause any overt tissue damage. To investigate if the differences in types and duration of nematode migration inside host tissue result in distinct plant growth responses, we analyzed changes in growth rate of the green canopy area at one day intervals following inoculation with 7.5 nematodes per gram of dry sand. We selected this density to represent a moderately severe infection which reduces plant growth without immediately overwhelming the plants (Fig 4a). This revealed three distinct patterns in the plant growth responses by nematode species. In the first few days after inoculation (up to 7 dpi), both *P. penetrans* and *H. schachtii* significantly reduced green canopy area growth, whereas *M. incognita* had no significant effect on green canopy area growth relative to mock-inoculated plants. Thereafter, when *P. penetrans* most likely alternated intracellular migration in the cortex with feeding periods, we observed a stimulation of green canopy growth of Arabidopsis plants. By contrast, from 8 to 19 dpi, when most infective juveniles of *H. schachtii* and *M. incognita* must have become sedentary, we observed a reduced growth of the green canopy area. After 19 dpi, the green canopy growth rates of plants inoculated with all three nematode species converged to a level slightly above that of mock-inoculated plants. Altogether our observations suggest that intracellular migration by nematodes has a strongly negative impact on plant growth but only during the onset of parasitism. Thereafter, the withdrawal of assimilates from the sap stream by feeding sedentary nematodes and the disruption of vascular connectivity by permanent feeding structures become more impactful on overall plant performance.

**Figure 4:**
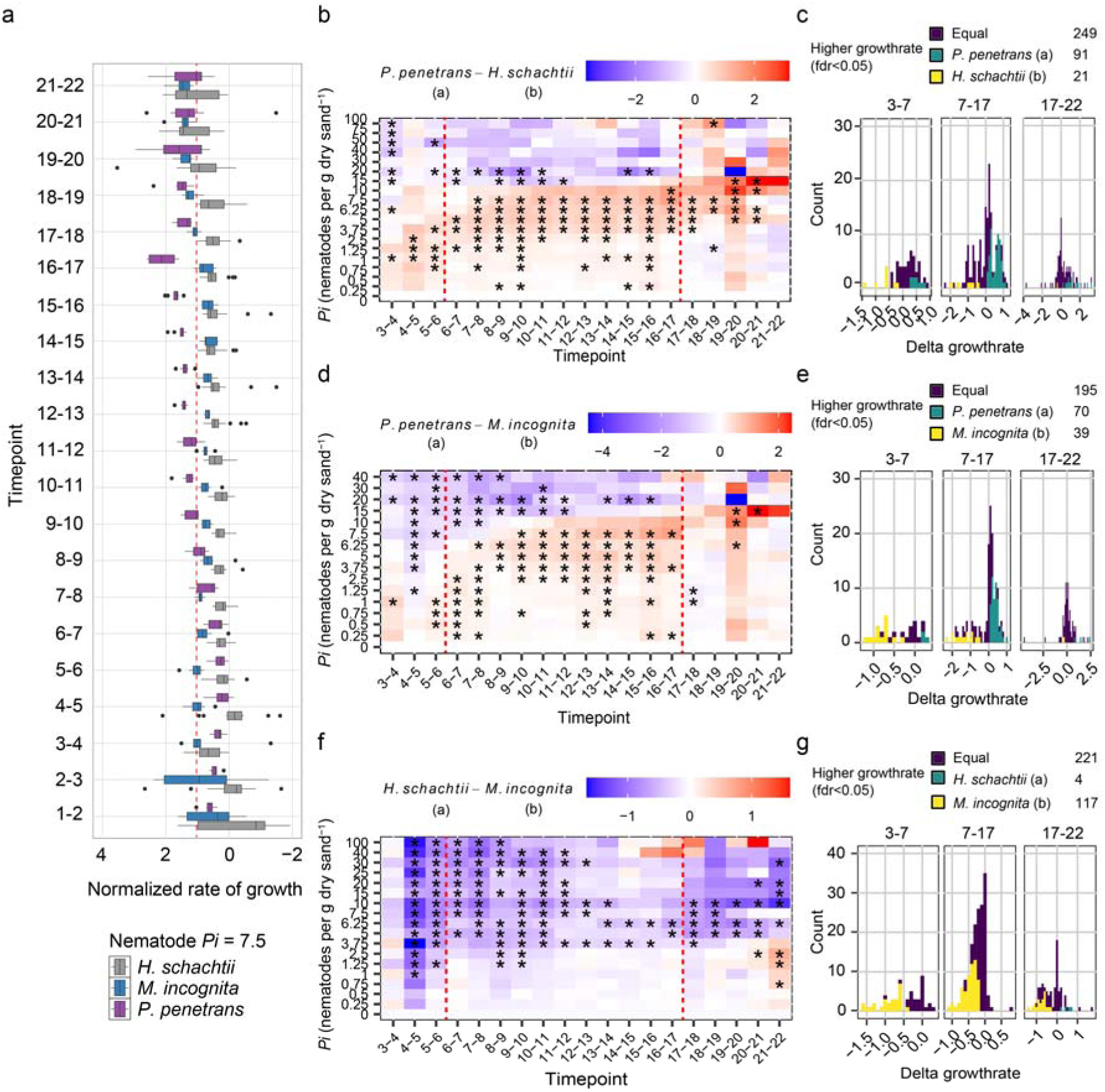
Comparison of growth rates of Col-0 inoculated with *P. penetrans*, *H. schachtii*, and *M. incognita*. Arabidopsis seedlings inoculated with 18 densities (*P_i_*) of *P. penetrans*, *H. schachtii*, or *M. incognita* (0 to 100 juveniles per g dry sand). Growth rates of green canopy areas were calculated from 1 to 22 dpi. **a)** Daily growth rates of plants treated with *P_i_* 7.5 were normalized to the corresponding growth rate of mock-treated plants. Red line indicates normalized growth rate of mock-inoculated plants. **b, d, f)** Heatmap of the differential normalized growth rate per *P_i_* and timepoint. Growth rates were normalized by dividing growth rates of individual plant by the median growth rate of mock treated plants (*P_i_* = 0). The difference in growth rate was calculated by subtracting normalized growth rates of nematode (a) inoculated plants from nematode (b) inoculated plants per timepoint and *P_i_*. Red indicates that plants inoculated with nematode (a) had a higher normalized growth rate than plants inoculated with nematode (b), and blue vice versa. Red dashed line indicates regions of interest for panels c, e, and g. **c, e, g)** Histograms of the number of occurrences that growth rates of nematode inoculated plants significantly differ per *P_i_* and timepoint. Differences in normalized growth rates between plants were tested using a paired t-test comparing data from the same timepoint and density combination, *p<0,05 (n=10-24). *H. schachtii* and *M. incognita* data originate from (Willig et al., 2023).

To more precisely pinpoint how Arabidopsis responds to stress induced by tissue damage and sedentarism, we performed pairwise comparisons of canopy growth rates across nematode densities and time intervals after inoculation (Fig. 4b, d, f). At relatively high inoculum densities (*P_i_* ≥ 10), *P. penetrans* reduced growth more strongly than both *H. schachtii* (Fig. 4b) and *M. incognita* (Fig. 4c) during the early phase of infection (<8 dpi). However, the differences between *P. penetrans* and the two sedentary species were broadly similar across the infection cycle (Fig. 4b and d). This is reflected in the distribution of favourable days, where *P. penetrans* more often showed higher growth rates than *H. schachtii* or *M. incognita* treated plants (Fig. 4c and e). In contrast, a pronounced difference was visible between the plant response to the two sedentary nematode species. While *H. schachtii* consistently reduced canopy growth, *M. incognita* had only a minor impact on plant growth, resulting in many more favourable days for *M. incognita* compared with *H. schachtii* (Fig. 4f and g). Thus, the strongest contrast in plant growth responses occurred between the two sedentary nematode species rather than between the migratory and two sedentary endoparasites.

### The formation of adventitious lateral roots is specific for cyst nematodes

Previously, we showed that Arabidopsis forms adventitious lateral roots in response to damage by *H. schachtii* in the first few days after inoculation (Guarneri et al., 2023; Willig et al., 2024). We therefore expected similar adaptations to root system architecture to occur in response to *P. penetrans*, which also causes clear tissue damage, but not to *M. incognita*. To test this hypothesis, we inoculated Arabidopsis seedlings *in vitro* with increasing numbers of *P. penetrans*, *M. incognita*, or *H. schachtii*, and counted the number of secondary roots at 7 dpi (Fig. 5a and b). Surprisingly, we did not observe an increase in the number of secondary roots following the inoculation with *P. penetrans*. Consistent with our previous observations, Arabidopsis seedlings inoculated with *H. schachtii* showed an increase in secondary roots by inoculation density. As expected, inoculation with *M. incognita* did not induce secondary root formation in Arabidopsis. To factor in differences in susceptibility of Arabidopsis for the three nematode species, we also counted the number of nematodes inside the root system (Fig. S3). At 7 dpi, the number of *P. penetrans* and *H. schachtii* inside the roots was not significantly different, while significantly fewer *M. incognita* had invaded Arabidopsis. This implies that the absence of additional secondary root formation upon inoculation with *P. penetrans* as compared to *H. schachtii* cannot be explained by a significantly smaller number of nematodes inside the root system. The induction of adventitious lateral roots is therefore specific for damage caused by *H. schachtii*.

**Figure 5:**
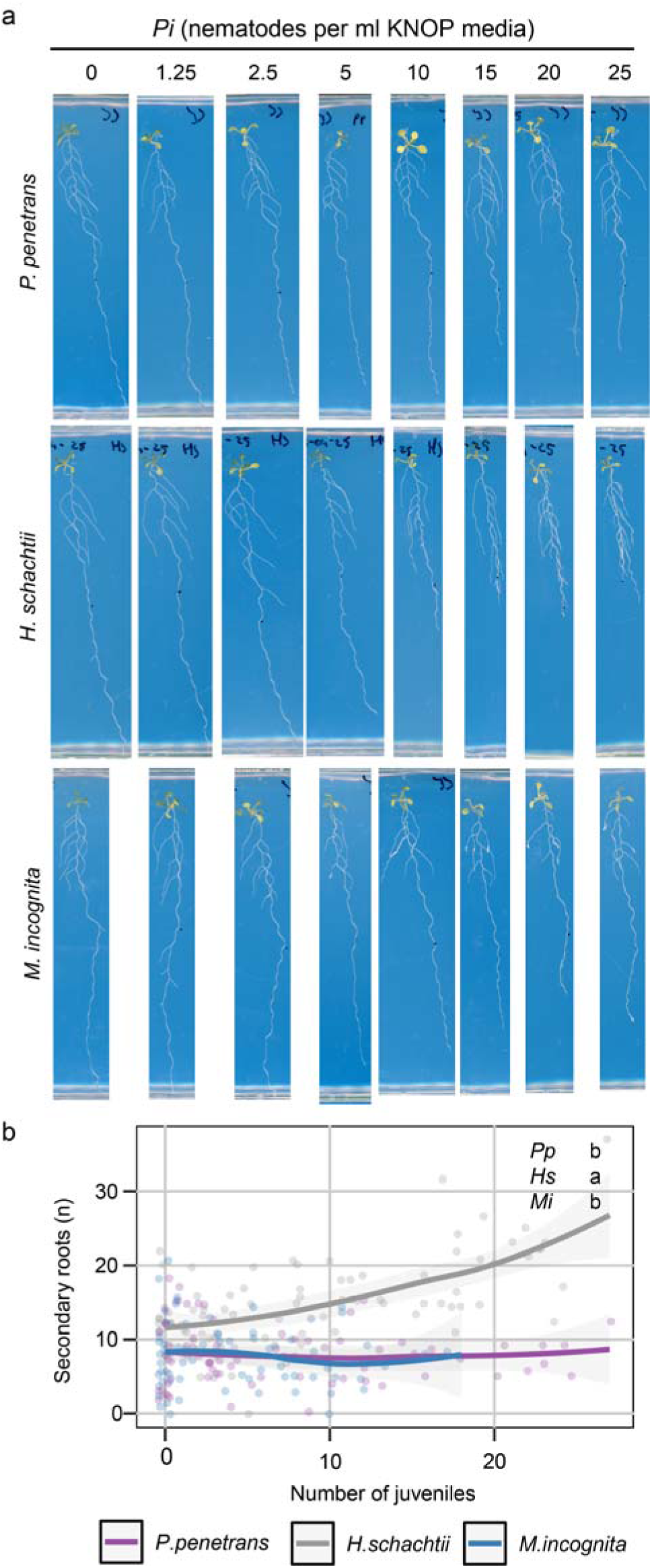
Adventitious lateral root formation is solely triggered by cyst nematode infection. Nine days old Arabidopsis seedlings (wild-type Col-0 or *arf7-1/arf19-1*) were inoculated with *P. penetrans*, *H. schachtii*, and *M. incognita*. At 7 dpi, scans were made of the root systems, and the number of secondary roots per plant or the number of plants forming secondary roots were counted. **a)** Representative pictures of wild-type Col-0 plants that have been inoculated with increasing inoculation densities (*P_i_*), ranging from 0 (mock) to 25 juveniles per mL KNOP media. **b)** Secondary roots formed per number of nematodes parasitizing (ecto- or endo-parasitism). Level of significance is indicated by letters. Significance of differences, between nematodes was calculated by analysis of variance followed by Tukey’s HSD test for multiple comparisons (*n* = 10-15; threshold for significance *P* < 0.05. Gray area indicates the 95% CI.

### Root system architecture of Arabidopsis remains largely unaffected by root-lesion nematodes

The induction of adventitious lateral roots by *H. schachtii* compensates for the loss of primary root growth, which contributes to tolerance of Arabidopsis to this nematode species (Guarneri et al., 2023). To investigate whether Arabidopsis employs different root plasticity mechanisms in response to *P. penetrans* and *M. incognita*, we also analysed total root length, primary root length, total secondary root length, and average secondary root length at 7 dpi by the number of nematodes inside the roots for these species (Fig. 6). At the time of inoculation, the nine days old Arabidopsis seedlings only had a single primary root, which restricted the number of *M. incognita* entering the root system via the root tip to a maximum of 18 nematodes per plant. High inoculation densities resulted in more nematodes inside the root system for *H. schachtii* and *P. penetrans* (Fig. 6 and Fig. S3), but for a valid comparison of root system architectures we used a cut-off for the data at 18 nematodes per plant for all species.

**Figure 6:**
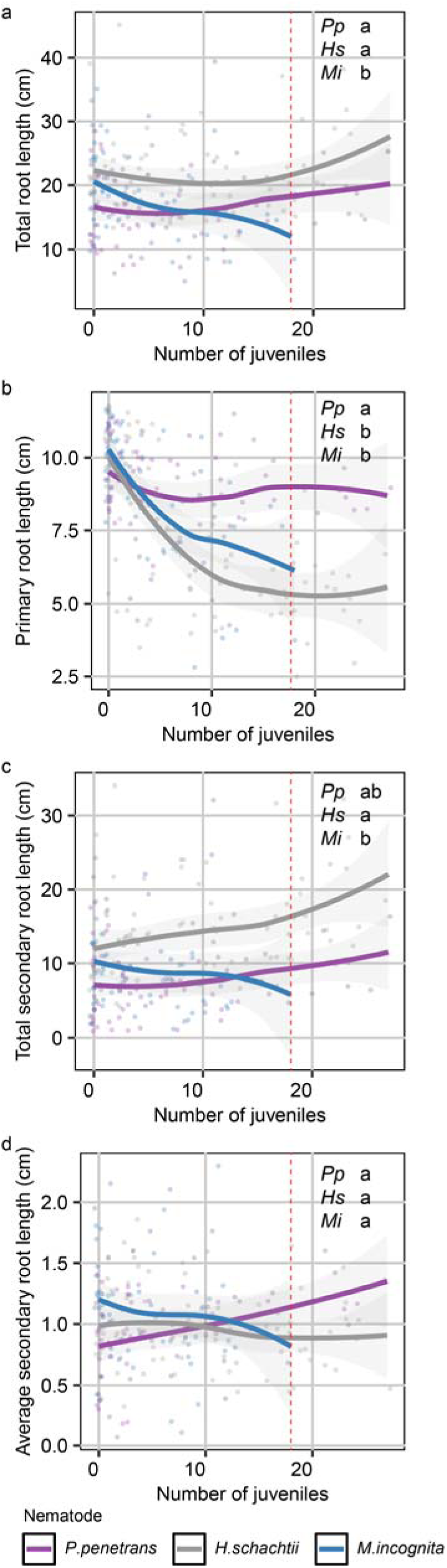
No compensatory root growth responses observed for *P. penetrans* and *M. incognita* infection in Arabidopsis. Arabidopsis seedlings (wild-type Col-0 or *arf7-1/arf19-1*) were inoculated with *P. penetrans*, *H. schachtii*, or *M. incognita*. At 7 dpi, scans were made of the root systems, and the root length was measured. **a)** Total root length that has grown per number of nematodes parasitizing. **b)** Primary root length that has grown per number of nematodes parasitizing. **c)** Total secondary root length that has grown per number of nematodes parasitizing. **d)** Average secondary root length that has grown per number of nematodes parasitizing. Level of significance is indicated by letters. Significance of differences between nematodes was calculated by analysis of variance followed by Tukey’s HSD test for multiple comparisons (*n* = 10-15; threshold for significance *P* < 0.05. Gray area indicates the 95% CI.

Interestingly, only *M. incognita* infections induced a significant decline in total root length by the number of nematodes inside the root (Fig. 6a). *M. incognita* also significantly reduced primary root length, similar to that of *H. schachtii*. Infections of *P. penetrans* did not affect primary root length (Fig. 6b). The total secondary root length slightly increased by the number of nematodes inside the root for *P. penetrans* and *H. schachtii*, which in the case of *P. penetrans* can be explained by a higher average secondary root length, while the number of secondary roots increased in the case of *H. schachtii* (Fig. 6c and d). *M. incognita* reduced the average secondary root length without inducing *de novo* secondary root formation resulting in the decline in total secondary root length. Based on these observations, we conclude that Arabidopsis compensates reduced primary root growth by *H. schachtii* by forming additional lateral roots, which increases the total secondary root length and stabilises the total root length. Arabidopsis does not compensate the impact of *M. incognita* on primary root growth by either forming additional secondary roots or increasing secondary root length resulting a decline of the total root length. Interestingly, *P. penetrans* induces a slight increase of the average secondary root length, which translates into a small increase in total secondary root length, but this does not have a significant effect on total root length.

## Discussion

Compensatory adaptations in root system architecture in response to local damage are thought to play an important role in plant tolerance to belowground biotic stress by root-feeding nematodes (Guarneri et al., 2023). This is based on the observation that the formation of additional adventitious lateral roots close to the feeding site of the sedentary endoparasite *H. schachtii* contributes to tolerance of Arabidopsis to this nematode species (Willig et al., 2024). At the outset of this study, it was not known if these observations could be generalised to nematodes with different invasion and feeding strategies. Here, we show that Arabidopsis tolerates migratory *P. penetrans* and sedentary *H. schachtii* equally well, while it is much more tolerant to sedentary *M. incognita*. These three nematode species differ in the level of damage they cause during migration, the location from where they draw their nutrients from the plant, and the impact of their feeding on host cells. In this paper we aim to link these differences in nematode biology to specific adaptations in Arabidopsis growth and development above- and belowground.

The tolerance limits of Arabidopsis for *P. penetrans* and *H. schachtii* are almost identical, but the underlying tolerance mechanisms are most likely not the same. Whereas Arabidopsis compensates for the impact of *H. schachtii* on primary root growth by forming additional secondary roots resulting in a stable total root length (Guarneri et al., 2023; Willig et al., 2024), *P. penetrans* does not significantly inhibit primary root growth and does not induce a compensatory response in root system architecture either. The absence of adaptations in root system architecture by increasing number of *P. penetrans* inside roots might explain the steep decline in plant performance and complete collapse of Arabidopsis above 15 nematodes per gram dry sand. Beyond this tipping point, the damage to the root cortex is probably so extensive that it is no longer able to facilitate movement of water and nutrients from surrounding growing soil to the xylem.

As the tolerance limit of Arabidopsis is much higher for *M. incognita* than for *H. schachtii* (Willig et al., 2023), we expected to find significantly more compensatory morphological adaptations in root system architecture for *M. incognita*. Surprisingly, although root-feeding by *M. incognita* has a major impact on primary root growth of Arabidopsis, we observed no formation of additional secondary roots or enhanced secondary root growth resulting in an overall decline in total root length. This implies that the tolerance of Arabidopsis for *M. incognita*, as opposed to *H. schachtii*, is not based on compensatory adaptations in root system architecture. As both *M. incognita* and *H. schachtii* employ sedentary feeding strategies, the basis for tolerance to *M. incognita* must most likely be found in differences in host invasion, location of feeding, or impact of feeding on host cells. The most obvious explanation is that *M. incognita* might simply causes less stress on plants and that this forms the basis of a higher tolerance limit (Guarneri et al., 2023; Zhang et al., 2019). But if that were true, we would expect to see similar root system adaptations to *M. incognita* as for *H. schachtii*, but at higher levels of infections. However, this is not in agreement with our observations.

A unique adaptation in root anatomy to *M. incognita* that might offer a better explanation for a higher tolerance limit is the formation of root knots or galls, made of host tissue, around giant cells. Root galls can be looked at as highly disordered anomalies in tissue organisation due to hormonal disruption by feeding nematodes, but gall formation may also be a tolerance mechanism to maintain the circulation within the vascular system of infected plants (Hoth et al., 2008; Olmo et al., 2020). In this sense, gall formation in host tissue surrounding giant cells may support the maintenance of vascular connectivity in the stele to secure the flow of assimilates, water, and nutrients. *De novo* vascularization which takes place inside galls (Bartlem et al., 2014) would then fit a dual purpose, i.e., supply feeding nematodes and maintain vascular connectivity. This hypothesis of gall formation as tolerance mechanism can be tested with Arabidopsis ecotypes which are less inclined to form galls in roots upon infection with *M. incognita*.

In our experiments, the reduced primary root growth was associated with feeding from adapted vascular cells by *M. incognita* and *H. schachtii*, which points at depletion of plant resources for the root apical meristem. As argued above, the formation of additional phloem vessels inside root galls may create a by-pass around the giant cells to maintain the acropetal flow of assimilates to the root tip, but this is evidently not enough to fully compensate for the extraction of assimilates by feeding root-knot nematodes. Likewise, the disruption of vascular connectivity and the extraction of assimilates from syncytia by feeding cyst nematodes may also deplete plant resources destined for the root apical meristem with evident consequences for primary root growth. However, *H. schachtii* induces formation of secondary roots close to infections sites (Guarneri et al., 2023; Willig et al., 2024), which will also compete with the primary root apical meristem for resources. Thus, the shorter primary roots observed in Arabidopsis seedlings infected with *H. schachtii*, as compared to *M. incognita*, may reflect a competition between the primary root apical meristem, feeding nematodes, and apical meristems of additional secondary roots.

We embarked on this study to investigate if different linages of nematodes can be used to investigate the molecular and cellular processes involved in mitigating the impact of different kinds of belowground biotic stresses in plants. We conclude that *P. penetrans* offers a model for studying plant responses to tissue damage specifically in the root cortex, focusing on the impact of breakdown of host cell integrity. In the first phase of parasitism, *H. schachtii* and *P. penetrans* have a similar impact on host tissue in the cortex, which implies that adventitious lateral root formation is not a compensatory mechanism to damage by nematode migration and feeding in the root cortex. Instead, the formation of additional secondary roots is more likely a specific response to nematode feeding from vascular cells, disruption of vascular connectivity, or both. Thus, infections by *H. schachtii* offer a model for studying the role of root system architecture plasticity in mitigating the impact of nematode feeding on differentiated vascular cells further away from the root apical meristem. By contrast, *M. incognita* offers a model to test hypotheses focusing on gall formation as a local tolerance mechanism to mitigate the impact of stress induced by nematode feeding close to the root tip.

## Supporting Information

Additional Supporting Information may be found in the online version of this article

## Author contributions

## Supplemental information

**Figure S1.**
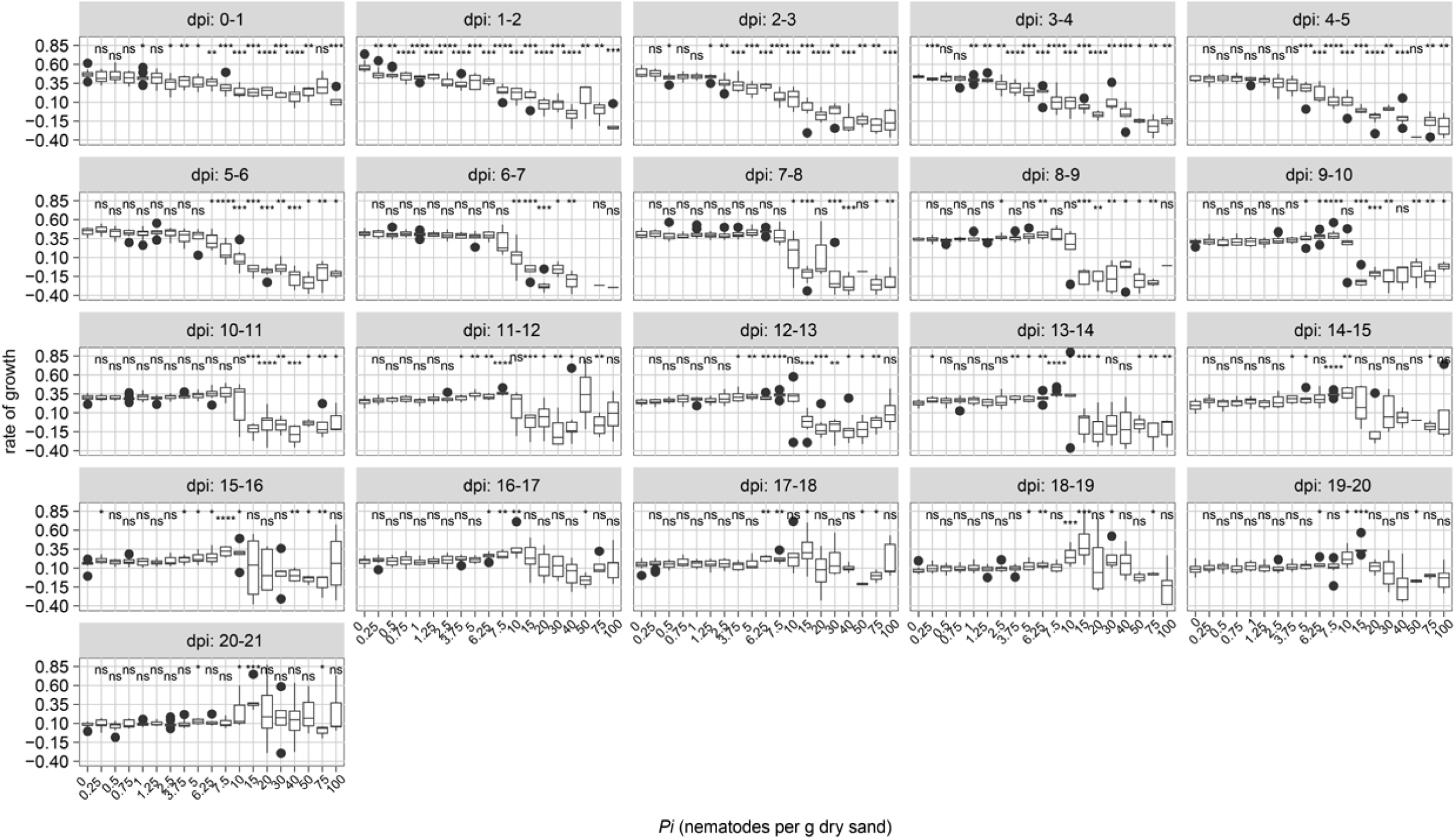
: Growth rate of Arabidopsis Col-0 plants inoculated with increasing densities of *Pratylenchus penetrans*. Nine-day-old Arabidopsis seedlings were inoculated with 20 densities (*P_i_*) of *P. penetrans* (0 to 100 nematodes per g dry sand). The growth rates of plants were calculated per day. Data was analysed with a Wilcoxon Rank Sum test. ns= not significant, *p< 0.05, **p< 0.01, ***p<0.0001 (n=10-24)

**Figure S2:**
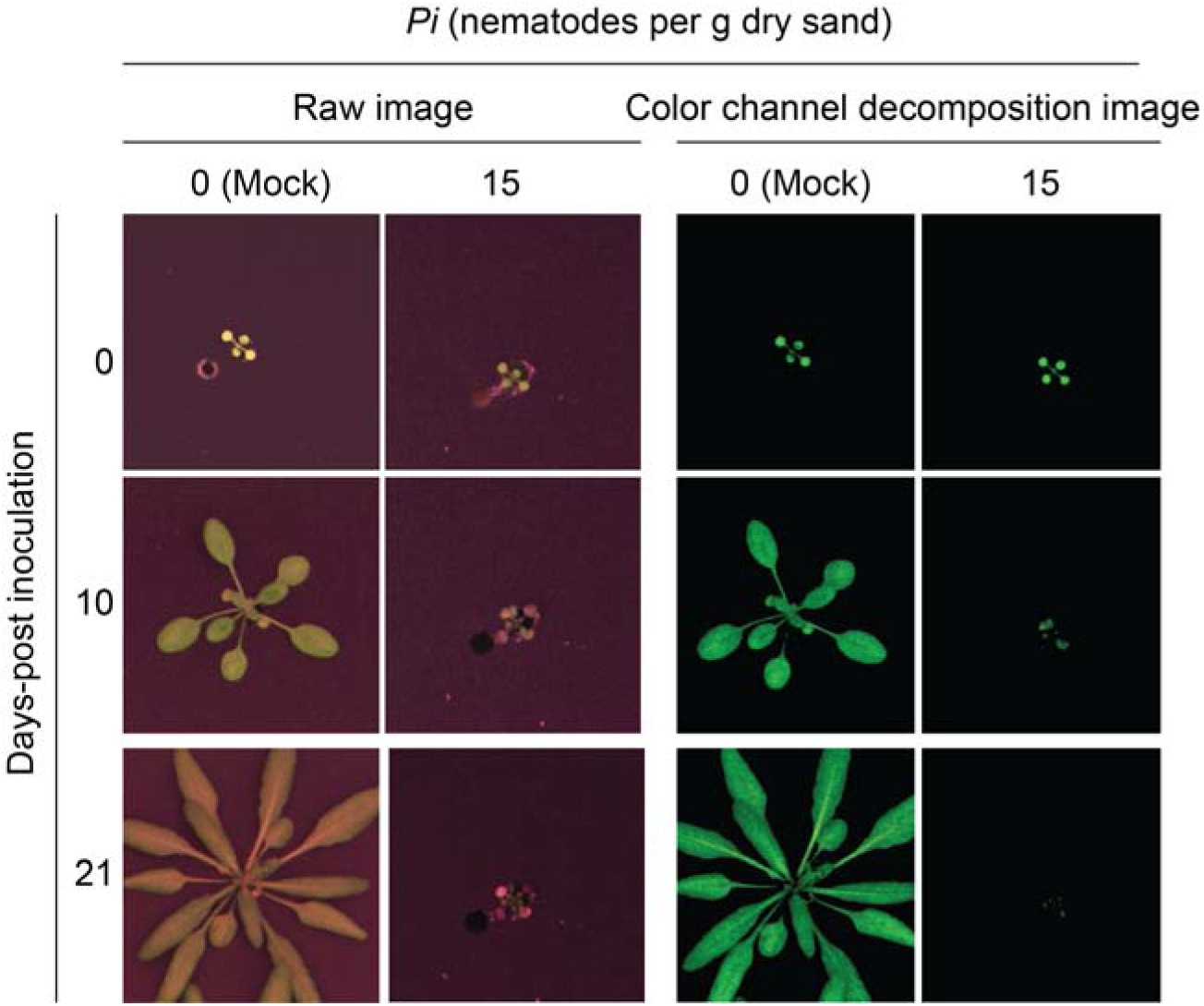
Col-0 seedlings treated with *Pratylenchus penetrans* inoculation density *P_i_*15 or higher completely collapsed at 10 dpi. Nine-day old Arabidopsis seedlings were inoculated with 20 densities (*P_i_*) of *P. penetrans* (0 to 100, *P. penetrans* (g dry sand)^-1^ and were monitored for a period of 21 days. Representative images were selected of plants inoculated with mock (0 *P. penetrans* juveniles) and *P_i_* 15 at 0, 10, and 21 days-post inoculation.

**Fig. S3:**
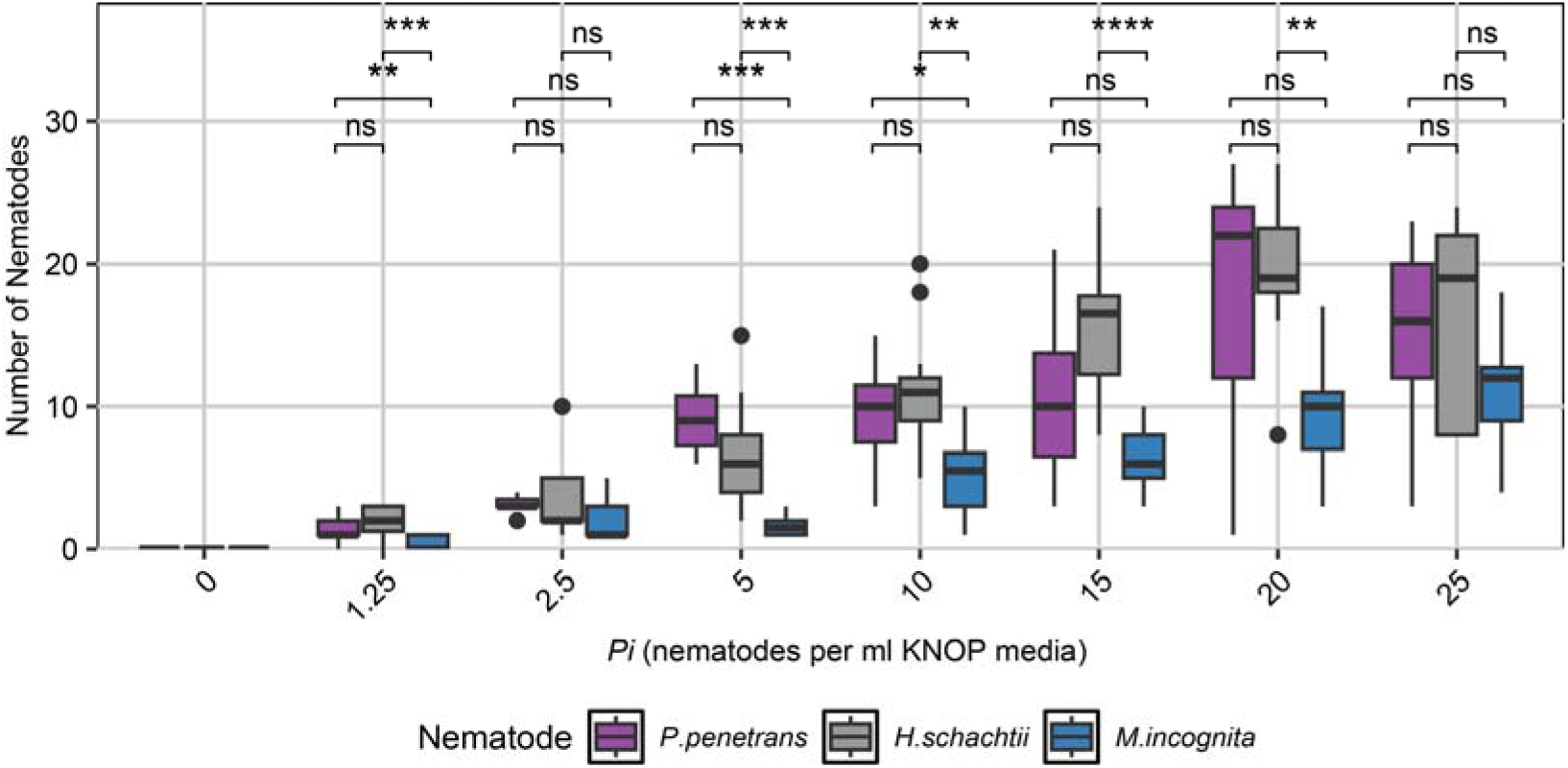
Arabidopsis Col-0 differs in its susceptibility to penetration by *P. penetrans*, *H. schachtii*, and *M. incognita*. Nine-day-old Arabidopsis seedlings were inoculated with nematode densities (*P_i_*) ranging from 0 (mock) to 25 juveniles (mL modified KNOP media)^-1^. Number of nematodes that successfully parasitized the root were counted after microscopy and fuchsin staining. Data from 2 independent biological repeats of the experiment were combined. The significance of differences between nematode species was calculated by ANOVA per *P_i_* (*n* = 10-15).

